# Identification of three sequence motifs in the transcription termination factor Sen1 that mediate direct interactions with Nrd1

**DOI:** 10.1101/492736

**Authors:** Yinglu Zhang, Yujin Chun, Stephen Buratowski, Liang Tong

## Abstract

The Nrd1-Nab3-Sen1 (NNS) complex carries out the transcription termination of noncoding RNAs (ncRNAs) in yeast, although the detailed interactions among its subunits remain obscure. Here we have identified three sequence motifs in Sen1 that mediate direct interactions with the RNA polymerase II CTD interaction domain (CID) of Nrd1, determined the crystal structures of these Nrd1 interaction motifs (NIMs) bound to the CID, and characterized the interactions *in vitro* and in yeast.

## Introduction

Nrd1 and Nab3 can form a complex through their dimerization domains and bind to RNA cooperatively through their RNA recognition modules (RRMs) (1–4) (Fig. 1a). The helicase Sen1 is speculated to remove nascent RNA from Pol II and trigger termination by collapsing the transcription bubble (5,6). Nrd1 also contains a CID that has a preference for Ser5P C-terminal domain (CTD) of Pol II (7,8), while Sen1 has a preference for Pol II Ser2P CTD (9). NNS couples tightly with the TRAMP complex for 3′-end trimming of ncRNAs and degradation of unstable RNAs (10–13), which is mediated by the interaction between Nrd1 CID and Trf4, a subunit of the TRAMP complex (14,15). The Nrd1 CID also interacts with Mpp6, a cofactor of the nuclear exosome (15). Besides ncRNA transcription termination (16), Sen1 also functions in R-loop resolution (17,18) and pre-mRNA transcription termination (19,20). However, how Sen1 interacts with Nrd1 and Nab3 to form the NNS complex is currently not well understood.

**Figure 1.**
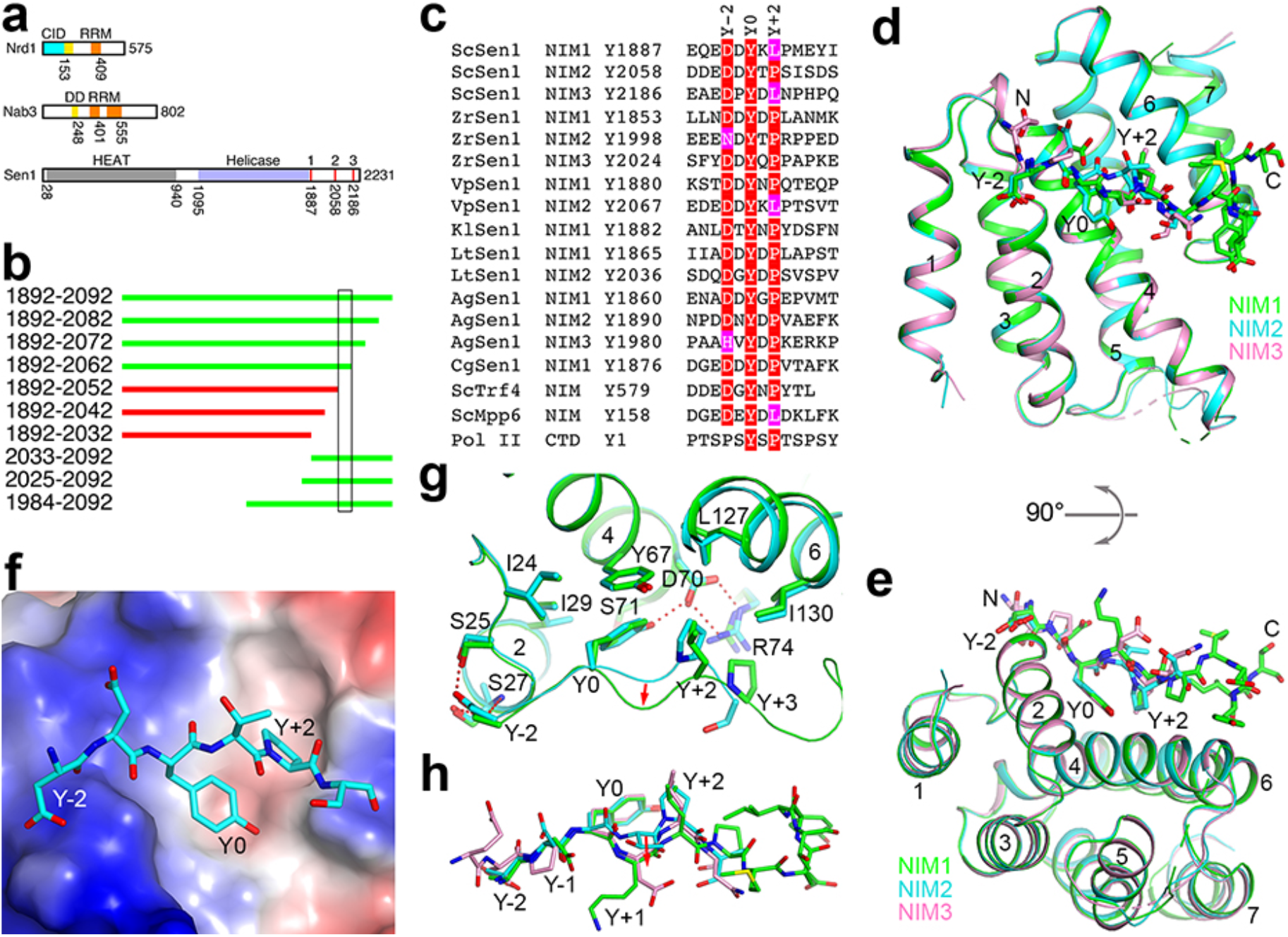
Identification of three NIMs in yeast Sen1. **(a)**. Domain organizations of yeast Nrd1, Nab3 and Sen1. The domains are given different colors, CID (cyan), DD (yellow), RRM (orange), HEAT repeats (gray), helicase (light blue). The three NIMs are in red. **(b)**. Various segments of Sen1 can (green) or cannot (red) pull down Nrd1 CID. The segment that is crucial for the interaction is boxed. **(c)**. Alignment of 15 NIMs found in seven fungal species. Also shown is the NIM in Trf4 and the Pol II CTD consensus. Residues in the consensus motif are highlighted in red and magenta. **(d)**. Overlay of the structures of CID in complex with NIM1 (green), NIM2 (cyan), and NIM3 (pink). **(e)**. Overlay of the structures, viewed after a rotation of 90° around the horizontal axis. **(f)**. Electrostatic surface of CID in the complex with NIM2 (stick models). **(g)**. Detailed interactions between NIM1 (green) and NIM2 (cyan) and the CID. Hydrogen-bonding interactions are indicated with the dashed lines (red). The conformational changes of the Y+1 and Y+2 residues are indicated with the red arrow. **(h)**. Overlay of the binding modes of the three NIM peptides. The structure figures were produced with PyMOL (www.pymol.org).

## Results and discussion

It was reported that the C-terminal segment (CTS) of *S. cerevisiae* Sen1 (ScSen1) (residues 1876-2231) interacts strongly with Nab3 (21). The CTS has few predicted secondary structures and no conservation among Sen1 homologs. However, we did not observe any interactions for purified *K. lactis* CTS with Nab3 and Nrd1 constructs that cover the region from the dimerization domain (DD) to the C-terminus (Figs. 1a, S1a). Surprisingly, we did observe a complex between a fragment of ScSen1 CTS (residues 1892-2092) and full-length Nrd1-Nab3 (Fig. S1b). Comparing the constructs used in these experiments, we hypothesized that the CID of Nrd1 mediated the formation of the complex. To confirm this hypothesis, we showed that deletion of the CID abolished complex formation (Fig. S1c). We then purified Nrd1 CID and demonstrated the direct interaction with purified Sen1 CTS by gel filtration (Fig. S1d). We also demonstrated interactions between the two proteins by pull-down experiments after co-expression in *E. coli* (Fig. S1e).

To map the region of ScSen1 CTS that is required for interaction with Nrd1 CID, we coexpressed various fragments of the CTS residues 1892-2092 with the CID, and discovered that Sen1 residues 1892-2062 could pull down CID but residues 1892-2052 could not (Figs. 1b, S1e, S1f). Residues 2052-2062 of Sen1 have the sequence DDDEDDYTPSI, and we realized that it is remarkably similar to the Nrd1 interaction motif (NIM) in Trf4 (Fig. 1c) (14,15), suggesting that ScSen1 also contains an NIM. The Tyr and Pro residues in Trf4 are important for binding the Nrd1 CID (14), and these residues are also present in the NIM of ScSen1.

A multiple sequence alignment of fungal Sen1 sequences aligned residues 2052-2062 of ScSen1 with DEDEDDYKLPT of *V. polyspora* Sen1 (VpSen1), suggesting that this could be another NIM. Surprisingly, we also found this motif in the CTS of ScSen1, near residue Tyr1887 (Fig. 1c) and just after the helicase domain (Fig. 1a), and confirmed its interaction with Nrd1 CID (Figs. S1g, S1h). Our crystal structure of its complex with the CID indicates that the Leu residue (rather than the Pro) is important for binding the CID (see below). Based on this information, we re-examined the ScSen1 CTS and found another sequence motif, near residue Tyr2186 (Fig. 1c), which we also confirmed by pull-down experiments (Fig. S1i). Therefore, ScSen1 has three NIMs in its CTS (Fig. 1a), and we have named them NIM1, NIM2, and NIM3 (Fig. 1c).

We then examined the CTS sequences of 12 fungal Sen1 and identified 15 NIMs in 7 fungal species, with a consensus motif (D/H/N)xYx(P/L), where x is frequently a hydrophilic residue (Fig. 1c). The Y-2 residue is predominantly Asp, while the Y+2 residue is predominantly Pro. Some species have three NIMs, while others have only one or two. Five of the fungal species examined, such as *C. albicans, D. hansenii* and *S. pombe*, do not appear to have an NIM in the CTS.

We have determined the crystal structures of Nrd1 CID in complex with each of the NIMs in ScSen1, at up to 2.0 Å resolution (Table S1). The structures of CID in the three complexes are essentially identical, with rms distance of 0.2-0.3 Å for equivalent Ca atoms between any pair of them (Figs. 1d, 1e). The NIMs bind at the same surface depression (Fig. 1f) as that for Pol II CTD (8) and Trf4 NIM (14,15) (Fig. S2). Clear electron density for the conserved (D/H/N)xYx(P/L) motif was observed in the complexes (Fig. S3). The remaining residues of the NIM peptides were mostly disordered, except the C-terminal residues of NIM1, which are stabilized by crystal packing. Therefore, residues outside of the conserved motif do not have strong interactions with the CID.

The NIM assumes a mostly extended conformation and has three major contacts with the CID, via the Y0, Y-2 and Y+2 residues (Fig. 1e). The Y0 residue of the NIM has a central role in the interaction with CID. Its side chain aromatic ring interacts with the side chains of Ile29, Tyr67 and the Y+2 residue, while its hydroxyl group is hydrogen-bonded to the side chain of Asp70, which is also involved in a bidentate interaction with Arg74 (Fig. 1g). The Y+2 residue contacts a hydrophobic patch formed by Leu127, Ile130, Tyr67 and the Y0 residue of the NIM, and the Asp70-Arg74 ion pair forms one side of the binding site for this residue. The Y-2 residue has hydrogen-bonding interactions with Ser25 and Ser27. It caps the N-terminus of helix a2, and the Asp side chain can also have favorable interactions with the dipole of this helix.

With Leu at the Y+2 position, a change in the backbone structure of the Y+1 and Y+2 residues is necessary to make room for this bulkier side chain compared to Pro (Figs. 1g, 1h). The binding modes of NIM1 and NIM3, which both have Leu at the Y+2 position, are similar.

Residues N-terminal to Y-2 are mostly negatively charged in ScSen1 as well as many other NIMs (Fig. 1c). An electro-positive patch in this region of Nrd1 CID (Fig. 1f) could suggest favorable interactions, but these residues in the NIMs are not ordered in the structures. *K. lactis* Sen1 has no negatively charged residues in this region, but it can interact with CID (Fig. S1h).

The structural observations are supported by our mutagenesis and fluorescence anisotropy binding assays. The *K*_d_ of the CID-NIM2 complex was 0.56±0.05 μM (Fig. 2a), similar to that of the CID-Trf4 NIM complex (14,15). In comparison, the *K*_d_ for the Y2058F mutant of NIM2 was 30-fold higher, confirming the importance of the Y0 residue for CID binding. Similarly, mutation of the Y-2 residue, D2056N, led to a 13-fold increase in the *K*_d_, while mutations of the Y+2 residue, P2060A and P2060V, led to ~20-fold increase in the *K*_d_ (Fig. 2b).

**Figure 2.**
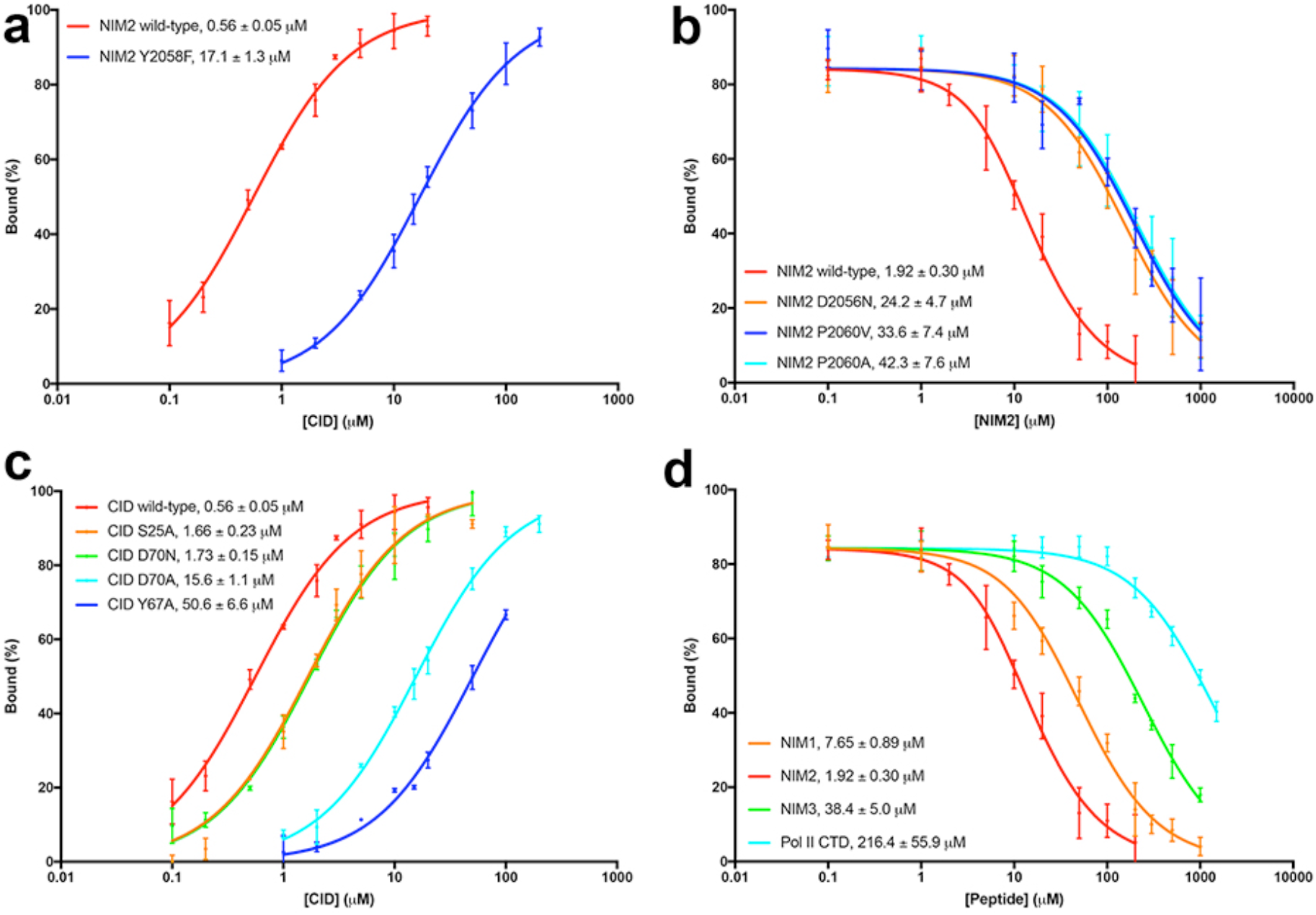
Binding affinity between Nrd1 CID and Sen1 NIMs. **(a)**. Fluorescence anisotropy binding data for wild-type NIM2 and the Y2058F mutant. The observed *K*_d_ values are indicated. **(b)**. Fluorescence anisotropy competition binding data for wild-type NIM2 and three mutants. **(c)**. Fluorescence anisotropy binding data for NIM2 with wild-type and mutant CID. **(d)**. Fluorescence anisotropy competition binding data for NIM1, NIM2, NIM3 and the Pol II Ser5P CTD. Error bars represent standard deviations from triplicate measurements.

We also introduced mutations in the NIM binding site of Nrd1 CID and characterized their effects on the interaction. The Y67A mutant showed a 90-fold increase in the *K*_d_ for NIM2, and the D70A mutant had a 30-fold increase (Fig. 2c). In comparison, the D70N mutant showed only 3-fold higher *K*_d_, indicating that the charge on this residue is not essential for NIM binding.

The *K*_d_ of the CID-NIM1 complex (7.6 μM) was 4-fold higher than that of the NIM2 complex, while the *K*_d_ of the CID-NIM3 complex (38 μM) was 20-fold higher (Fig. 2d). The *K*_d_ of the Mpp6 peptide, which also has a Leu at the Y+2 position (Fig. 1c), is 13.6 μM (15). Together with the data on the P2060A and P2060V mutants, our observations suggest that several hydrophobic residues could be tolerated at the Y+2 position. In comparison, the Pol II Ser5P CTD peptide studied here had the weakest affinity for the CID, with a *K*_d_ of 216±56 μM, comparable to the value reported previously (14). The binding mode of the CTD peptide has substantial differences to the NIMs (Fig. S2d). The CTD also contains multiple repeats, which could provide increased avidity for the interaction with Nrd1.

We showed that mutation of the Tyr residue in all three NIMs in Sen1 CTS (Y1887A, Y2058A, Y2186A) was required to abolish the interaction with CID, while single-site mutants were still able to interact with CID (Fig. S4). We also confirmed that other regions of Sen1, such as its helicase and HEAT repeat domains, could not interact with Nrd1 CID, and that Nrd1 lacking the CID could not mediate the formation of the NNS complex (Fig. S5). Overall, these data indicate that, under the conditions tested here, Sen1 interacts with Nrd1 solely through the NIM-CID contacts, and that Sen1 does not interact with Nab3.

We next characterized the Sen1-Nrd1 interaction in yeast. Deletion of the CTS had no effect on growth (Fig. S6), consistent with earlier data (22), nor did mutating the NIM tyrosines to alanine, individually or in combination (data not shown). A Nrd1 mutant lacking the CID also has no growth or transcription termination defects (7), indicating that none of the Nrd1 CID interactions are essential under laboratory growth conditions. On the other hand, removing or mutating NIMs from the Sen1 CTS had a profound effect on the interaction with Nrd1, based on co-immunoprecipitation experiments (Fig. 3). A Sen1 mutant lacking all three NIMs showed essentially no binding to Nrd1. A mutant with one NIM showed weaker binding compared to one with two NIMs, and mutation of the Tyr residue to Ala in this NIM essentially blocked binding to Nrd1.

**Figure 3.**
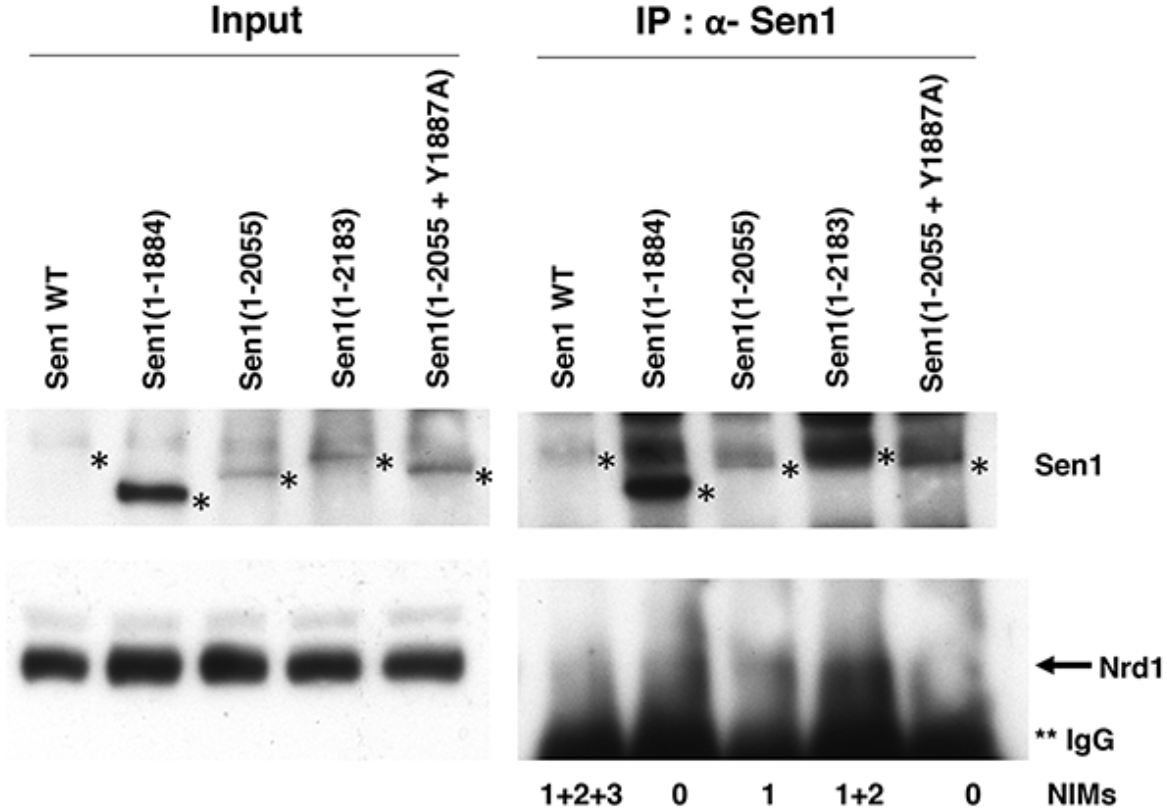
Sen1 NIMs are crucial for interaction with Nrd1 in yeast. Cell extracts from the indicated Sen1 strains were immunoprecipitated with anti-Sen1 antibody and probed for the presence of Nrd1 and Sen1 proteins. Positions of Sen1 proteins are marked with a single asterisk. Nrd1 is partially obscured by its proximity to IgG heavy chain on the blot, marked with two asterisks.

As we and others have previously noted (23), normal levels of full-length Sen1 are extremely low and difficult to detect by Western blotting. Correspondingly little Nrd1 was detected upon immunoprecipitation (Fig. 3). Certain N-terminal truncations stabilize Sen1 by removing degradation-promoting sequences (23,24). We found that the C-terminal truncations of Sen1 also produced higher levels of Sen1 protein, suggesting additional degradation signals in the CTS.

In conclusion, we have identified three NIMs in the CTS of budding yeast Sen1 and elucidated the molecular basis for their recognition by Nrd1 CID. The identification of one of these NIMs (NIM2) in ScSen1 was independently reported recently (25). Although neither the Sen1 NIMs nor Nrd1 CID is essential for supporting viability, their conservation suggest these interactions are very likely to promote NNS function. The Nrd1 CID now has four known binding partners, Sen1, Trf4 (14), Mpp6 (15) and Pol II CTD (7,8), and other motifs that bind this CID could also be possible. In fact, the crystal structure of free Nrd1 CID (7) contains a packing contact that places a YLNA motif, from helix a7 that has partially unwound, in the binding site, with the Ala residue occupying the Y+2 position (Fig. S3e). Further studies will shed light on whether additional motifs could actually support interactions between Nrd1 CID and other proteins, and how the Sen1-Nrd1 interactions mediate the functions of the NNS complex.

## Supporting information

## Acknowledgments

This research is supported by NIH grants R35GM118093 and S10OD012018 (to LT) and R01GM056663 (to SB). We thank Dave Brow (University of Wisconsin, Madison) for Sen1 and Nrd1 materials.

## Materials and Methods

### Protein expression and purification

*S. cerevisiae* Nrd1 CID (residues 6-151) was subcloned into the pET28a vector (Novagen). The recombinant protein, with an N-terminal hexa-histidine tag, was over-expressed in *E. coli* BL21 Star (DE3) cells (Novagen), which were induced with 0.4 mM IPTG and allowed to grow at 16 °C for 16–20 h. The soluble protein was purified by nickel-charged immobilized-metal affinity chromatography, ion-exchange chromatography and gel filtration chromatography. The purified protein was concentrated in a buffer containing 20 mM Tris (pH 7.0), 150 mM NaCl, 10 mM DTT, and stored at -80 °C. The His-tag was not removed prior to crystallization.

### Protein crystallization

Crystals of Nrd1 CID-Sen1 NIM complex were grown at 20 °C with the hanging-drop vapor diffusion method. Nrd1 CID protein solution was at 20 mg/ml concentration, and the protein was mixed with Sen1 NIM peptides (GL Biochem) at a molar ratio of 1:10. The reservoir solution contained 0.1 M sodium citrate (pH 4.0), 1 M lithium chloride and 20% (w/v) PEG 6000. Fully-grown crystals were obtained one day after set-up. The crystals were cryo-protected in the crystallization solution supplemented with 15% (v/v) ethylene glycol and flash-frozen in liquid nitrogen for data collection at 100 K.

### Data collection and processing

X-ray diffraction data sets were collected at a wavelength of 0.979 Å on a Pilatus-6MF pixel array detector at the 24-ID-C beamline of the Advanced Photon Source (APS). The diffraction images were processed and scaled with XDS (26).

### Structure determination and refinement

The structures were solved by molecular replacement with the program Phaser-MR in PHENIX (27,28). The crystal structure of *S. cerevisiae* Nrd1 CID (PDB code 3CLJ) (7) was used as the search model. Manual model rebuilding was carried out with Coot (29). The structure refinement was performed with the program PHENIX, with translation, libration, and screw-rotation (TLS) parameters. The data processing and refinement statistics are summarized in Table S1.

### Fluorescence anisotropy assay

Nrd1 CID mutants were expressed and purified using the same protocol as that for the wild-type CID. NIM1, NIM2, NIM3, NIM2 mutants, Pol II Ser5p CTD and N-terminally 5-carboxyfluorescein (5-FAM) labeled NIM2 peptides were obtained for fluorescence anisotropy (GL Biochem). For direct binding assays, 5 nM 5-FAM labeled NIM2 peptide was titrated with wild-type and mutants of Nrd1 CID at increasing concentrations. For competition assays, complex of 5 nM 5-FAM labeled NIM2 peptide and 3 μM wild-type Nrd1-CID was titrated with unlabeled NIM1, NIM2, NIM3, NIM2 mutants and Pol II Ser5p CTD peptides at increasing concentrations. The measurements were conducted on a Synergy Neo2 Hybrid Multi-Mode Reader (Biotek). Samples were excited at 485 nm and emissions were recorded at 528 nm. Measurements were performed in triplicates at room temperature in 20 mM Tris (pH 7.0), 200 mM NaCl, 5 mM DTT. The binding curves were fitted by GraphPad Prism (GraphPad Software, La Jolla) using hyperbolic equation for direct binding and the analytical model for competitive binding (30).

### Mixing assay

Mixtures of purified proteins were incubated on ice for 1 h and separated on Superose 6 Increase 10/300 GL column (GE Healthcare) in 20 mM Tris (pH 7.0), 200 mM NaCl, 5 mM DTT.

### Pull-down assay

Various regions of *S. cerevisiae* Sen1 CTS and its mutants and *K. lactis* Sen1 CTS were sub-cloned into the pET28a-SUMO vector. The recombinant Sen1 proteins would contain an N-terminal hexa-histidine tag followed by a SUMO tag. The plasmid was cotransformed with one for un-tagged *S. cerevisiae* Nrd1 CID (residues 6-151) or *K. lactis* Nrd1 CID (residues 1-155) and over-expressed in *E. coli* BL21 Star (DE3) cells (Novagen). The culture was induced with 0.4 mM IPTG and allowed to grow at 16 °C for 16–20 h. The soluble protein was incubated with nickel-charged resin for 1 h at 4 °C, washed with a buffer containing 20 mM Tris (pH 7.0), 200 mM NaCl, 20 mM imidazole and eluted with a buffer containing 20 mM Tris (pH 7.0), 200 mM NaCl, 150 mM imidazole. The eluate was mixed with protein dye and run on an SDS-PAGE gel.

### Yeast genetics

All yeast strains are derived by plasmid shuffling into YSB3181 (ura3Δ0, leu2Δ0, trp1Δ::LEU2/KanR, his3Δ1, met15Δ0, sen1Δ::KanMX, [pRS416 + -700 Sen1]) using selection on selective media containing 5-fluoroorotic acid. For growth assay shown in Fig. S6, serial dilutions of post-selection cells were spotted on YPD plates. The following strains carry the listed plasmids expressing the indicated Sen1 mutants:

1. YSB3514: Sen1 pRS414 + -700 Sen1 (WT)
2. YSB3515: pRS313-Sen1 (WT) (22)
3. YSB3516: pRS313-Sen1(1-1858) (22)
4. YSB3517: pRS414-Sen1(1-1884)
5. YSB3518: pRS313-Sen1(1-1907) (22)
6. YSB3519: pRS414-Sen1(1-2055)
7. YSB3520: pRS414-Sen1(1-2183)
8. YSB3521: pRS414-Sen1-Y1887A YSB3522: pRS414-Sen1(1-2053)
9. YSB3523: pRS414-Sen1(1-2055 + Y1887A)

### Immunoprecipitation experiments

Overnight cultures of yeast were inoculated into 50 ml YPD at OD595 of 0.3 and grown for about five hours to OD 1.0 at 30 °C. Cells were pelleted, washed, and re-suspended in nucleosome isolation buffer (0.1% Triton X-100, 10 mM MgCl_2_, 20 mM HEPES (pH 7.8), 250 mM sucrose) plus 150 mM NaCl, and protease (leupeptin, pepstatin A, PMSF, aprotinin) and phosphatase inhibitors (sodium fluoride and sodium vanadate) (31). Cells were lysed by vortexing with glass beads for six 30 second intervals with cooling on ice in between. Debris were removed by microfuging at 13000 rpm for 20 minutes at 4 °C. After decanting the supernatant, 10 μl of 10 mg/ml RNaseA was added to make sure any interactions were not mediated by RNA. For immunoprecipitations, 2 mg of extracts were incubated with 2 μl of anti-Sen1 (a gift from Dave Brow, University of Wisconsin) and 20 μl of Protein A-Sepharose beads overnight at 4 °C. Beads were gently pelleted and washed three times for five minutes with the same buffer used during immunoprecipitation.

Bound proteins were eluted into loading buffer, separated by SDS-polyacrylamide gel electrophoresis, and blotted at 45 volts to nitrocellulose overnight at 4 °C. After blocking, membranes were probed with anti-Sen1 or Anti-Nrd1 (also provided by Dave Brow), using antirabbit IgG-HRP conjugate and ThermoFisher Pico Plus developer to image bands on X-ray film.

