## Supplementary material for "Identification of three sequence motifs in the transcription termination factor Sen1 that mediate direct interactions with Nrd1"

**Table S1**
**Data collection and refinement statistics**

| | CID+NIM1 | CID+NIM2 | CID+NIM3 |
|---|---|---|---|
| **Data collection** | | | |
| Space group | $C222_1$ | $C222_1$ | $C222_1$ |
| Cell dimensions | | | |
| a, b, c (Å) | 103.4, 109.1, 86.7 | 98.9, 102.5, 115.5 | 98.7, 103.0, 115.4 |
| α, β, γ (°) | 90, 90, 90 | 90, 90, 90 | 90, 90, 90 |
| Resolution range (Å)[1] | 50-2.1 (2.2-2.1) | 50-2.0 (2.1-2.0) | 50-2.8 (3.0-2.8) |
| No. of observations | 185,949 | 175,380 | 48,822 |
| No. of unique reflections | 27,909 | 39,419 | 14,439 |
| $R_{merge}$ (%) | 8.4 (67.9) | 8.0 (48.7) | 14.7 (48.8) |
| I/σI | 16.8 (5.4) | 12.2 (3.1) | 7.3 (2.8) |
| $CC_{1/2}$ | | | |
| Completeness (%) | 96.2 (96.7) | 98.1 (96.9) | 97.1 (97.2) |
| Redundancy | 6.7 (6.9) | 4.4 (4.3) | 3.4 (3.4) |
| **Refinement** | | | |
| Resolution range (Å) | 46-2.1 (2.2-2.1) | 45-2.0 (2.1-2.0) | 45-2.8 (2.9-2.8) |
| No. of reflections | 27,890 | 39,414 | 14,426 |
| $R_{work}$ (%) | 18.6 (23.0) | 17.9 (25.2) | 23.7 (29.1) |
| $R_{free}$ (%) | 20.4 (28.6) | 22.6 (29.2) | 30.0 (31.3) |
| No. atoms | | | |
| Protein | 2,428 | 3,561 | 3,560 |
| Cl | 0 | 2 | 2 |
| Water | 126 | 261 | – |
| *B*-factors | | | |
| Protein | 42.7 | 35.4 | 40.9 |
| Cl | – | 23.5 | 31.2 |
| Water | 45.0 | 40.2 | – |
| RMS deviations | | | |
| Bond lengths (Å) | 0.006 | 0.007 | 0.008 |
| Bond angles (°) | 0.73 | 0.76 | 1.10 |
| Ramachandran plot statistics (%) | | | |
| Most favored region | 98.29 | 98.85 | 97.92 |
| Additional allowed region | 1.37 | 1.15 | 2.08 |

[1]The numbers in parentheses are for the highest resolution shell.

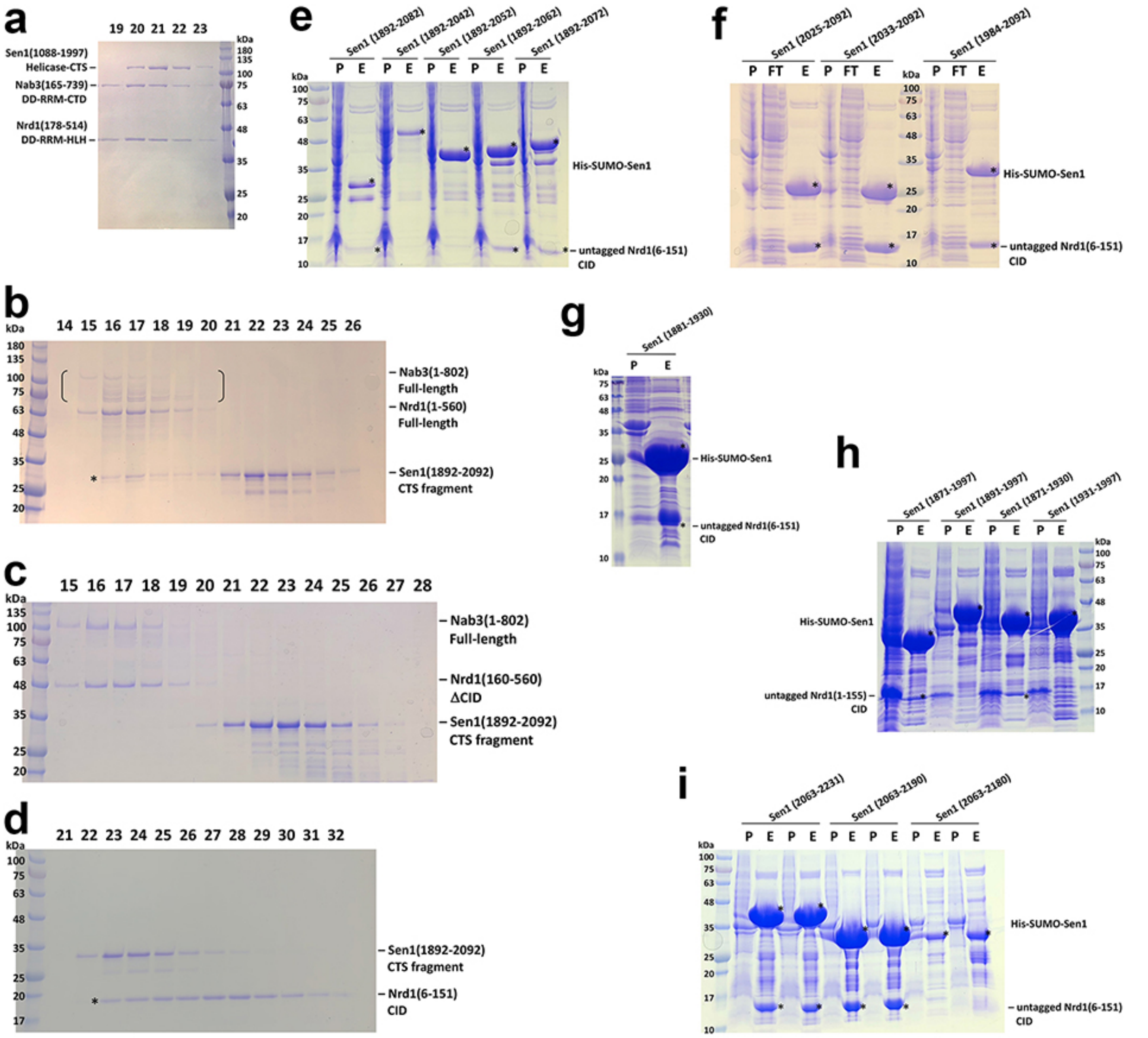

**Fig. S1. Identification of three segments in Sen1 that interact with Nrd1 CID. (a-d)**. Mixtures of purified Sen1, Nrd1 and Nab3 proteins were run on a gel filtration column, and the indicated fractions were resolved by SDS-PAGE. **(e-i)**. His-tagged Sen1 were co-expressed with untagged Nrd1 CID to map their regions of interaction. P: pellet, E: Ni column eluate, FT: flow through. Panels a and h are for *K. lactis* proteins, and all other panels for *S. cerevisiae* proteins.

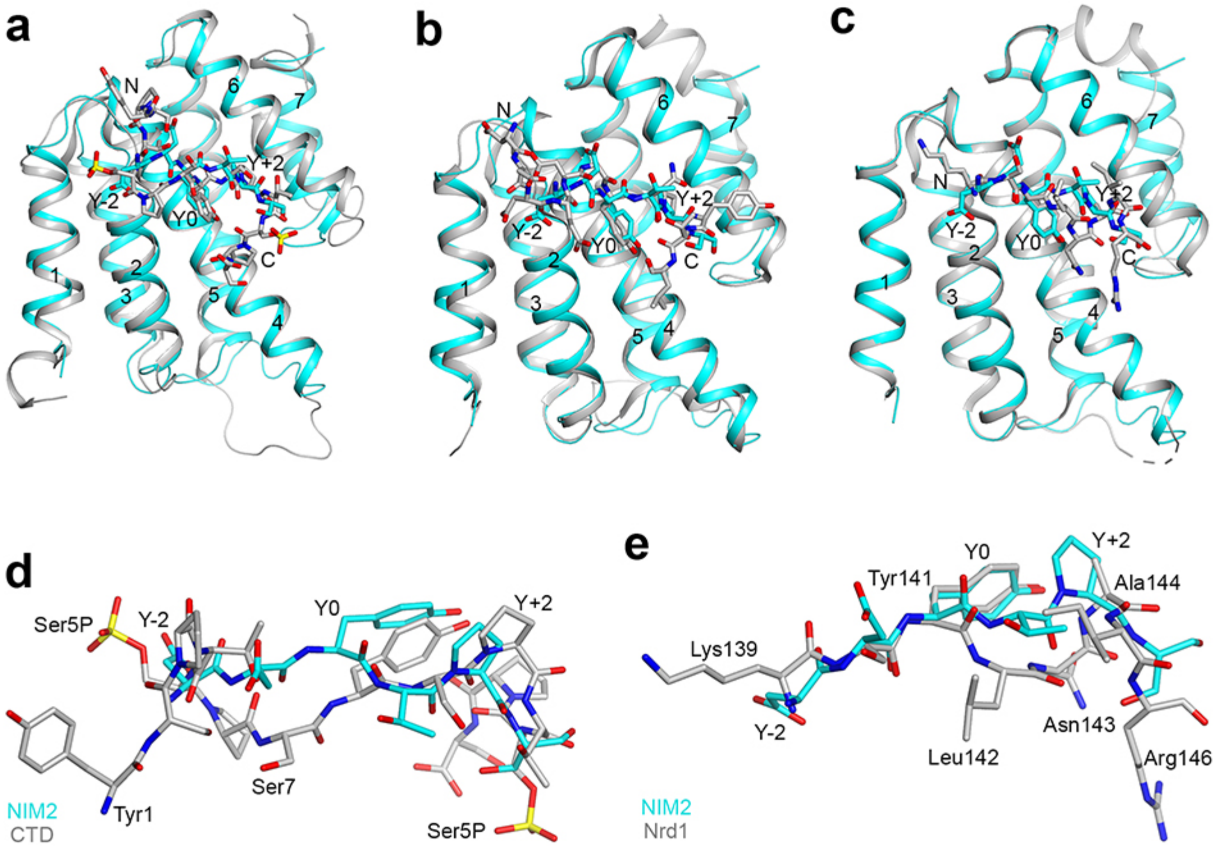


**Fig. S2. Comparison of the binding modes of other sequence motifs to Nrd1 CID. (a)**. Overlay of the NIM2 complex (cyan) with the Ser5P CTD complex (gray). **(b)**. Overlay of the NIM2 complex (cyan) with the Trf4 NIM complex (gray). **(c)**. Overlay of the NIM2 complex (cyan) with the Nrd1 CID in complex with residues 139-146 from another molecule by crystal packing (gray). These residues are in helix α7, which has become partially unwound. **(d)**. Overlay of the binding modes of NIM2 (cyan) with the Ser5P CTD (gray). **(e)**. Overlay of the binding modes of NIM2 (cyan) with the Nrd1 CID residues 139-146 from crystal packing (gray).

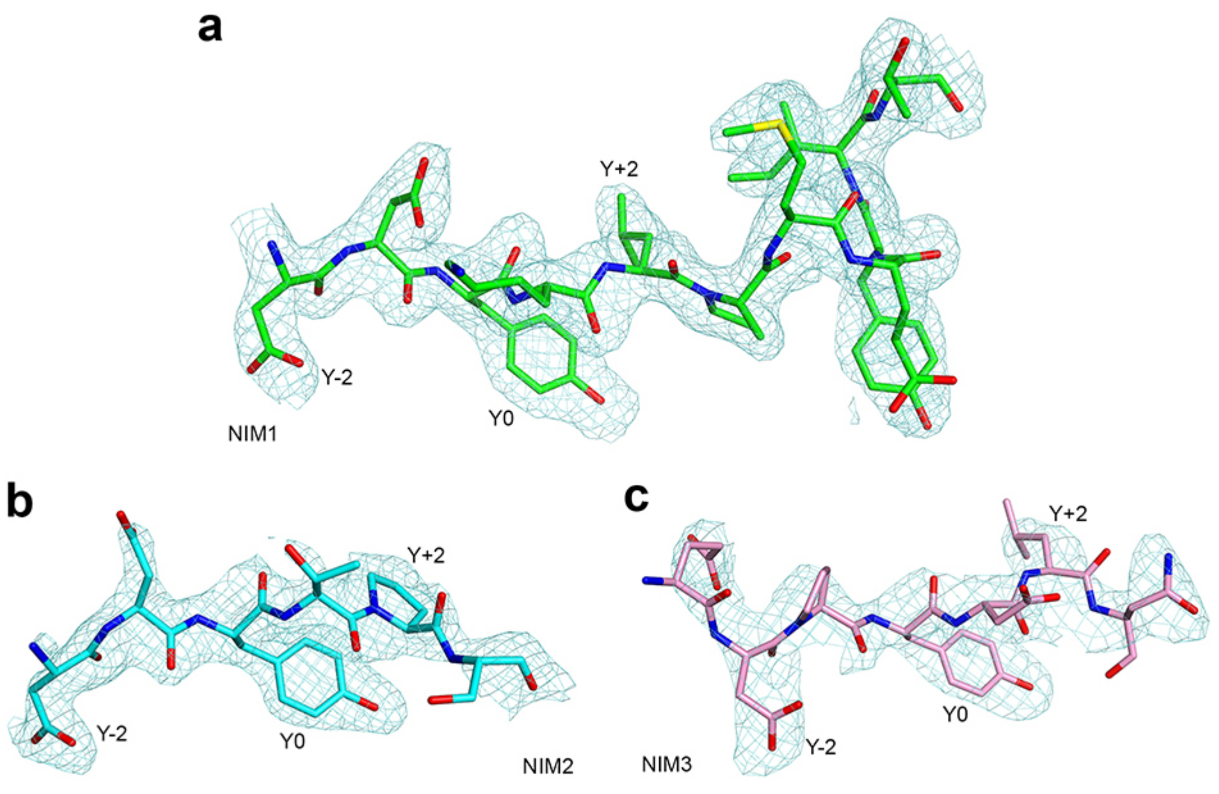


**Fig. S3. Observed electron density for the NIMs.** Omit Fo–Fc electron density map for **(a)** NIM1 at 2.1 Å resolution, contoured at 2.5σ; **(b)** NIM2 at 2.0 Å resolution, contoured at 2.5σ; **(c)** NIM3 at 2.8 Å resolution, contoured at 2σ.

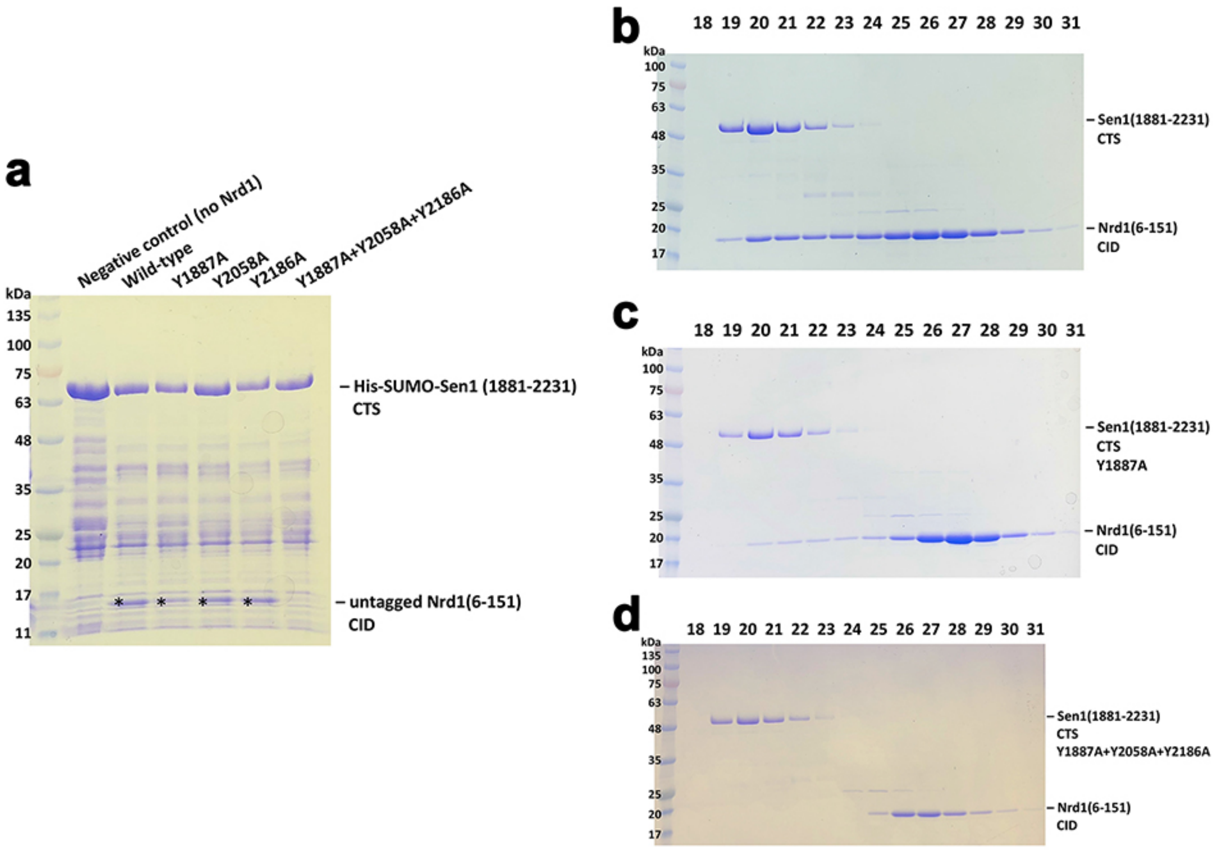


**Fig. S4. Mutation of all three NIMs is necessary to block Sen1-Nrd1 interaction. (a)**. His-tagged Sen1 CTS wild-type and various NIM mutants were co-expressed with untagged Nrd1 CID. The Ni column eluate was then resolved on SDS-PAGE. **(b-d)**. Mixing experiments with purified Sen1 CTS (wild-type and indicated mutants) with Nrd1 CID.

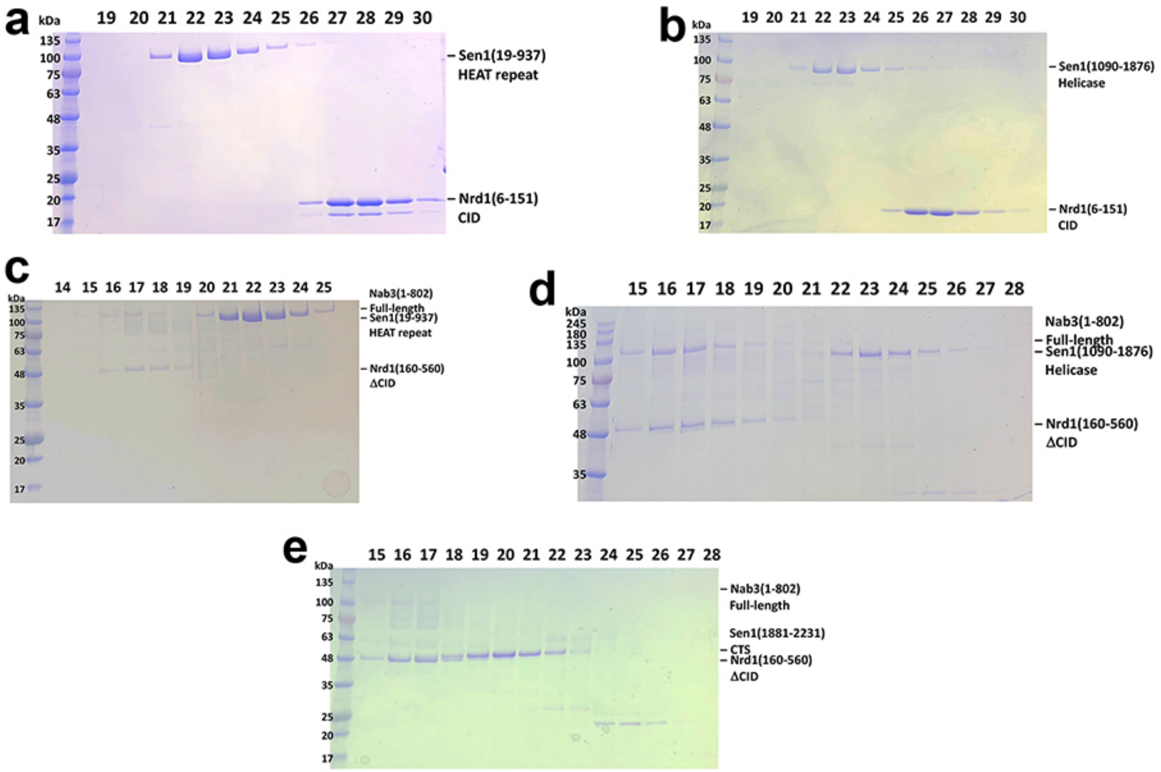


**Fig. S5. Sen1 interacts with Nrd1 only through the CID.** Mixtures of purified Sen1, Nrd1 and Nab3 proteins were run on a gel filtration column, and the indicated fractions were resolved by SDS-PAGE. **(a)**. No interaction between Sen1 HEAT repeat domain and Nrd1 CID. **(b)**. No interaction between Sen1 helicase domain and CID. **(c)**. No ternary complex among full-length Nab3, Sen1 HEAT repeat domain, and Nrd1 lacking the CID. **(d)**. No ternary complex among full-length Nab3, Sen1 helicase domain, and Nrd1 lacking the CID. **(e)**. No ternary complex among full-length Nab3, Sen1 CTS, and Nrd1 lacking the CID.

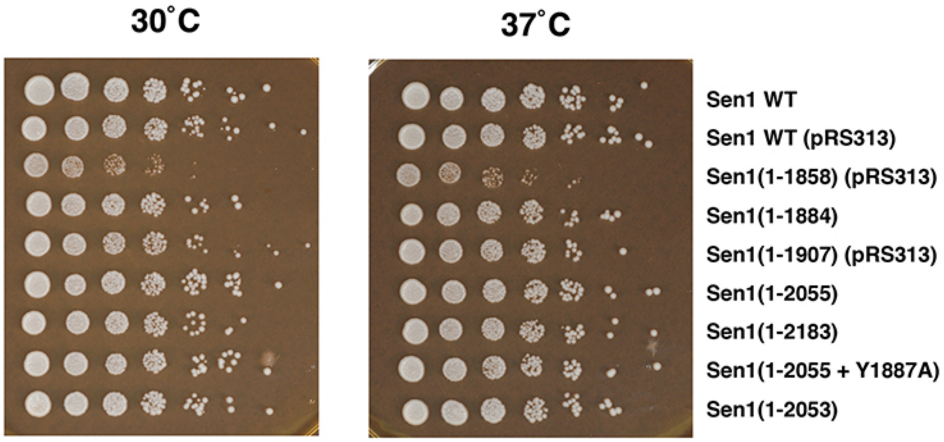


**Fig. S6. Sen1 CTS is not required for growth**. The indicated Sen1 mutants were created by plasmid shuffling and tested for growth by serial dilution spotting assay on rich YPD media. The Sen1(1-1858) mutant is missing the C-terminal region of the helicase domain, which may explain its growth phenotype.
